# MNase profiling of promoter chromatin in *S. typhimurium*-stimulated GM12878 cells reveals dynamic and response-specific nucleosome architecture

**DOI:** 10.1101/816348

**Authors:** Lauren Cole, Jonathan Dennis

**Affiliations:** Department of Biological Science, Florida State University, Tallahassee, FL 32306

**Keywords:** chromatin, nucleosome, micrococcal nuclease, GM12878, *Salmonella typhimurium*

## Abstract

The nucleosome is the primary unit of chromatin structure and commonly imputed as a regulator of nuclear events, although the exact mechanisms remain unclear. Recent studies have shown that certain nucleosomes can have different sensitivities to micrococcal nuclease (MNase) digestion, resulting in the release of populations of nucleosomes dependent on the concentration of MNase. Mapping MNase sensitivity of nucleosomes at transcription start sites genome-wide reveals an important functional nucleosome organization that correlates with gene expression levels and transcription factor binding. In order to understand nucleosome distribution and sensitivity dynamics during a robust genome response, we mapped nucleosome position and sensitivity using multiple concentrations of MNase. We use the innate immune response as a model system to understand chromatin-mediated regulation. Herein we demonstrate that stimulation of a human lymphoblastoid cell line (GM12878) with heat-killed *Salmonella typhimurium* (HKST) results in widespread nucleosome remodeling of response-specific loci. We further show that the response alters the sensitivity of promoter nucleosomes. Finally, we correlate the increased sensitivity with response-specific transcription factor binding. These results indicate that nucleosome distribution and sensitivity dynamics are integral to appropriate cellular response and pave the way for further studies that will deepen our understanding of the specificity of genome response.

## Introduction

The functional role of chromatin is inseparable from an appropriate cellular response to a stimulus. The fundamental subunit of chromatin is the nucleosome, composed of a histone octamer core and approximately 150bp of DNA (Kornberg, 1999). It is commonly asserted that the occupancy and position of nucleosomes can provide access to the underlying DNA sequence, thus affecting nuclear processes (Kaplan et al., 2009). The distribution of nucleosomes across the genome is controlled by chromatin remodeling complexes and DNA sequence (Gupta et al., 2008). However, characterization of the functional organization of the genome remains a major challenge in biology today. Micrococcal nuclease (MNase) was first used to isolate nucleosomal DNA from the chicken beta-globin gene (Sun et al., 1986). It remains the predominant method for generation of nucleosome occupancy maps in eukaryotic genomes, and previous studies have mapped changes in chromatin structure during differentiation, environmental stimulus or stress, and disease states (Shivaswamy et al., 2008; Teif et al., 2012; Druliner et al., 2013; Sexton et al., 2014, Sexton et al., 2016). Additionally, it has been recently shown that nucleosomes exhibit differential sensitivity to MNase. The sensitivity of promoter nucleosomes, particularly the +1 and −1 nucleosomes relative to the transcription start- and regulatory factor-binding sites, is a defining chromatin characteristic that gives insight into chromatin-mediated regulation of these loci. The importance of characterizing this differential sensitivity of chromatin has been demonstrated in multiple organisms and is an important variable to consider (Vera et al. 2014, Mieczkowski et al., 2016, Cherije et al. 2019).

In order to investigate the role of nucleosome dynamics during the innate immune response, we have mapped nucleosome position and sensitivity at all human promoters in GM12878 lymphoblastoid cells stimulated with heat killed *Salmonella typhimurium* (HKST). We mapped nucleosome distributions with two MNase levels: one heavy digestion level and one light digestion level. The comparison of these different MNase digestion levels reveals important information about transcription factor binding, gene expression prediction, and sensitivity of chromatin to digestion (Vera et al., 2014, Mieczkowski et al., 2016, Pass et al., 2017, Brahma and Henikoff 2019). The complicated interplay between chromatin remodeling complexes and the specific epigenetic landscape is largely unknown and is likely responsible for the diverse kinetics of the immune response (Ramirez-Carrozzi et al., 2006, Natoli 2010, Dorrington and Fraser 2019). Here we show that stimulation of a human lymphoblastoid cell line with HKST causes changes in promoter nucleosome distribution as well as changes in sensitivity to MNase. We find that 10% of promoters in the 20m HKST treatment gain nucleosome occupancy upstream of the TSS (−1 nucleosome) compared to the untreated control. Furthermore, we find that the −1 nucleosome becomes less sensitive to MNase during the 20m HKST treatment and that immune transcription factor binding at the TSS (NFkB, Pu1, and Ebf1) is associated with flanking nucleosomes sensitive to MNase digestion.

## Materials and Methods

### Cell culture

Cell GM12878 cells were grown at 37°C in 15% FBS-supplemented RPMI medium. Cells were stimulated with 1.0×10^9^ heat-killed Salmonella typhimurium (HKST; 15 minutes at 80°C) for 20 minutes, 40 minutes and 60 minutes and harvested at the end of each time point.

### Cell harvest and nuclei purification

Approximately 1 × 10^7^ cells were harvested, cross-linked in 1% formaldehyde, and incubated for 10 min at room temperature. After the 10 min incubation, the cross-linking reaction was quenched with 125 mM glycine. Next, the nuclei were isolated in nucleus isolation buffer containing: 10 mM HEPES at pH 7.8, 2 mM MgOAc_2_, 0.3 M sucrose, 1 mM CaCl_2_, and 1% Nonidet P-40. The nuclei were then pelleted by centrifugation at 1000g for 5 min at 4°C.

### MNase digestion of chromatin

At each time point ~2.5×10^6^ nuclei were treated with light (40U MNase/mL) and heavy MNase-digestion conditions (400U MNase/mL), see average fragment size distribution in **Fig. S1B**. The chromatin at each time point was digested with the respective concentration of MNase for five minutes at 37°C and stopped with EDTA. DNA from MNase-digested nuclei was isolated via phenol-chloroform extraction and mononucleosome sized bands resolved with a 2% agarose gel. The ~150bp mononucleosomal band was excised for each time point and MNase concentration. Additionally, two untreated control samples were harvested and processed as described, and have been referenced here as untreated samples.

### Mononucleosome DNA Library Preparation

MNase sequencing libraries were prepared using NEBNext Ultra DNA library Prep Kit for Illumina (NEB #E7370S), using 30ng of input mononucleosomal DNA from each digestion level and time point. Following end-prep and adaptor ligation, the libraries were purified with AMPure XP beads. Universal and index primers from NEBNext Multiplex Oligos for Illumina (NEB #E7335S) were incorporated by a 12 cycle PCR. Library size and quality was verified with the Agilent 2100 Bioanalyzer. Molar concentration of indexed library was determined by KAPA quantitative PCR and size corrected using sizing information from the Bioanalyzer.

### Solution-based Sequence Capture and Illumina Flowcell hybridization and sequencing

Previously, we combined MNase-seq with in-solution targeted enrichment of 2 kb surrounding TSSs of 21,857 human genes (Druliner et al., 2016; Sexton et al., 2016), as curated by NCBI RefSeq (Pruitt et al., 2009). We termed this approach Transcription Start Site MNase-seq (mTSS-seq). Size selected fragments (~50-200 bp) were used to prepare Illumina sequencing libraries and subjected to targeted enrichment utilizing the custom-designed Roche Nimblegen SeqCap EZ Library. DNA fragments were captured according to the Roche Nimblegen protocol. By qPCR we observe ~300 fold enrichment of sample target genes compared to off-target loci. Paired-end reads (see below) were aligned to the hg19 genome assembly (IHGSC 2001) and read densities were inspected with the UCSC genome browser.

### HiSeq 2500 data processing

Illumina adapters were clipped and aligned to the HG19 genome assembly, with unpaired and non-uniquely aligned reads discarded (bowtie2, samtools). Mononucleosome-sized fragments were used to infer nucleosome position. Nucleosome occupancy profiles were obtained by calculating the fragments per million that mapped at each base-pair in the SeqCap regions (bedtoolsCoverage). Midpoints for nucleosome distributions were determined through the calculation of center fragments in 60 bp windows at a 10 bp step in the 2kb surrounding each TSS. Data was subsequently processed in R (https://github.com/dvera).

### Data Availability

The data discussed in this publication have been deposited in NCBI’s Gene Expression Omnibus (Edgar et al., 2002) and are accessible through GEO Series accession number GSE139224.

## Results

### HKST stimulation of B-lymphoblastoid cells results in changes in promoter nucleosome occupancy

We have mapped nucleosome distribution during the immune response to HKST. <u>M</u>Nase <u>T</u>ranscription <u>S</u>tart <u>S</u>ite-enriched sequencing (mTSS-seq) allows for high quality nucleosome maps at all human promoters (**Fig. S1**). mTSS-seq data is highly concordant with the preeminent nucleosome maps in lymphoblastoid cell lines (**Fig. S1C**, Gaffney et al., 2012). Work from our lab has measured widespread and transient nucleosome redistributions during viral reactivation (Sexton BS, 2014, Sexton BS, 2016). To determine whether nucleosome redistributions were limited to viral reactivation or a broader feature of the immune response, we mapped nucleosome distributions at 20, 40, and 60 minutes following stimulation with HKST. We observed the greatest differences in nucleosome distribution at 20 minutes after stimulation with HKST. We calculated the differences in nucleosome distribution between the untreated cells and the HKST-stimulated time points at all promoters measured. We observe that average nucleosome profile remains similar between the untreated control and timepoints (**Fig. 1A**). However, over a third of all human promoters fall into four distinct clusters which reflect gain or loss of nucleosome occupancy at the nucleosomes +1 and −1 relative to the TSS: documented regulatory nucleosomes (**Fig. 1B**) (Field et al., 2008; El Gazzar et al., 2010, Rando 2012). Cluster 5 in **Figure 1B**reflects a gain of occupancy at the −1 nucleosome, and those promoters are significantly enriched for immune processes and transcription (**Fig. 1C**, **Table S1**). Cluster 1 (**Figure 1B**) represents loci with more nucleosomal fragments in the untreated control cells, and gene ontology analysis for this cluster reveals significant enrichment for developmental processes, post-transcriptional silencing, and differentiation (Eden et al., 2009, **Table S2**). These results suggest that nucleosome occupancy changes over the course of HKST treatment. Specifically, 20 minutes after HKST stimulation, we observe that 2271 genes (~10% of promoters) show increased nucleosomal occupancy at the −1 position, and these genes are enriched for transcription regulation and immune-specific processes. Finally, 35% of the genes in cluster 5 (**Figure 1B**) are also in the top quartile of expressed genes in GM12878 cells. They are ontologically enriched for B-cell antigen processing and presentation of exogenous antigen and recombination of immunoglobulin genes involved in immune response, as well as many important growth-associated processes (**Table S1**).

**Figure 1.**
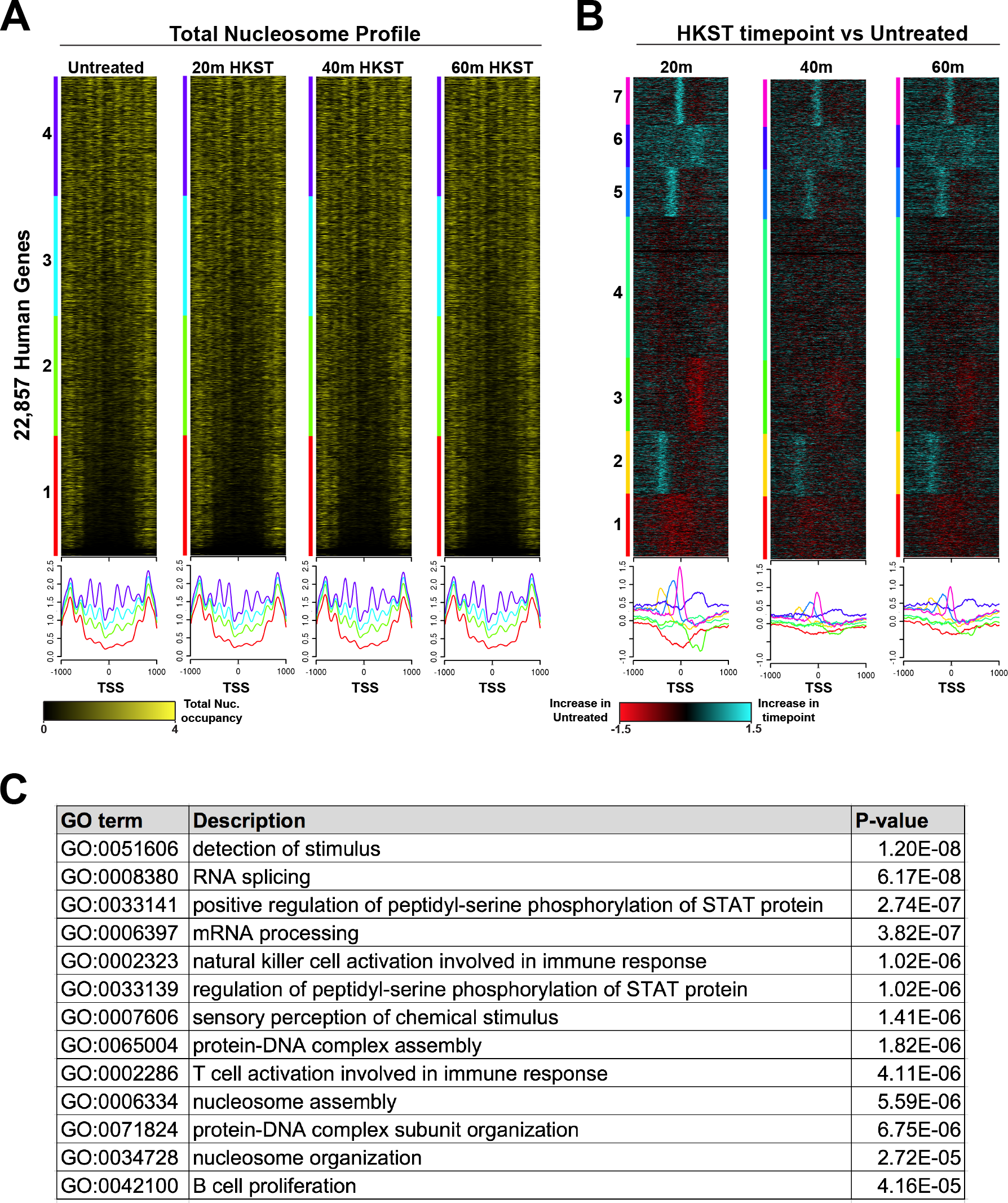
Nucleosome occupancy changes during time course stimulation with HKST. (A) Total nucleosome fragments sorted into quartiles based on maximum signal. All heatmaps sorted in the same order, –1000bp/+1000bp surrounding the TSS for all RefSeq open reading frames. Yellow indicates presence of nucleosomal fragment. (B) The log2ratio of the untreated sample and HKST time points: 20, 40, and 60 minutes post-stimulus. Resulting Log2ratio heatmaps sorted by kmeans clustering (k=7). All heatmaps sorted in the same order, –1000bp/+1000bp surrounding the TSS for all RefSeq open reading frames. Red indicates more nucleosomal fragments present in the untreated GM12878 cells and cyan indicates increased nucleosomal fragments in the stimulus timepoint. (C) Table of gene ontology enrichment for the promoters which gain–1 nucleosome fragments in the 20m HKST timepoint.

### HKST stimulation of B-lymphoblastoid cells results in changes in MNase sensitivity of promoter nucleosomes

We use differences in nuclease sensitivity to distinguish nucleosome-bound fragments that are readily released when lightly digested (MNase-sensitive fragments, MSFs) from nucleosome-bound fragments that are resistant to heavy digestion (MNase-resistant fragments, MRFs). To identify MSFs within our mTSS-seq experiment we calculated the log2ratio of the light/heavy MNase digests. We then sorted the resulting MNase-sensitivity nucleosome profiles based on maximum expression using existing RNAseq datasets for GM12878 cells (Davis et al., 2018). We also visualized active transcription using a ChIP-seq dataset of CTD phosphorylated RNA polymerase II (Pol2ser2) sorted on the same sort order (Davis et al., 2018). We observed astrongly positioned −1 sensitive nucleosome in the top quartile of maximum active transcription, and that this nucleosome becomes more sensitive in the 20m post-HKST timepoint (**Fig. 2A**). We also observe less positioned nucleosomes, containing a mix of sensitive and resistant nucleosomes in the bottom quartile of low and non-expressed genes (**Fig. 2A**). Consistent with a primary response to bacterial infection (Herschman 1991, Winkles 1998), the largest sensitivity differences were seen at the 20 minute timepoint (**Fig. S2**)

**Figure 2.**
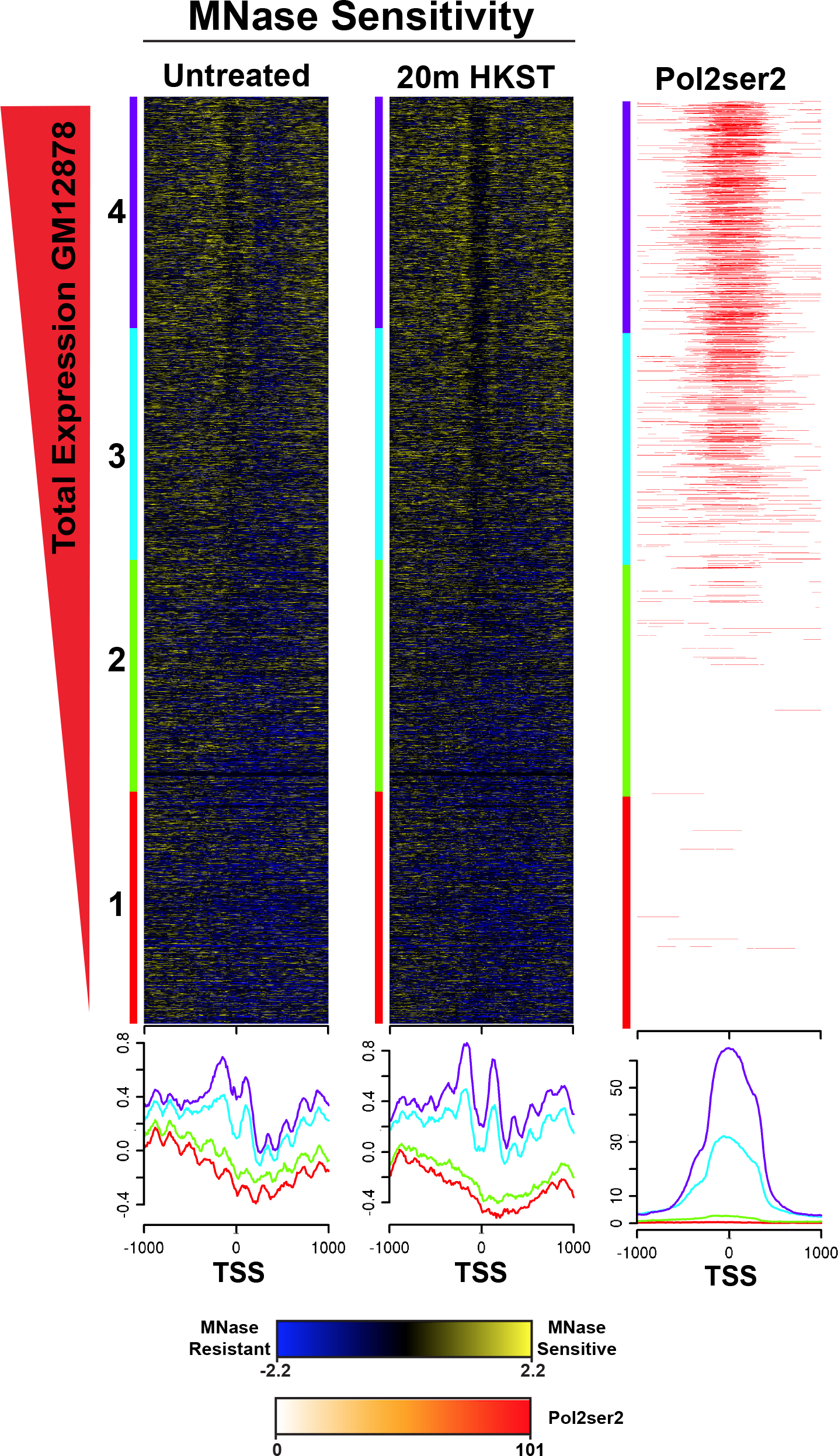
Sensitive nucleosomes are associated with transcription and nucleosome sensitivity changes during HKST stimulation. MNase sensitivity of untreated GM12878 and 20m HKST-treated cells were sorted into quartiles based on total expression (GSM2344230). ChIPseq data for actively transcribing Pol2ser2 was sorted into quartiles based on the same sort order. Blue indicates MNase-resistant nucleosomal fragments and yellow indicates MNase-sensitive nucleosomal fragments.

### Sensitivity of TSS-flanking nucleosomes is associated with transcription factor binding

We next wanted to determine the relationship between MNase-sensitivity and transcription factor binding. As we have associated nucleosome sensitivity with active transcription, we sorted the 20 minute HKST time point into quartiles based on maximum sensitivity surrounding the TSS. We then sorted transcription factor ChIP-seq data for NFkB, Pu1, and Efb1 in the same gene order (**Fig. 3**) (Davis et al., 2018). Here we show that MSFs flank important immune regulatory factors at the TSS during the immune response to HKST (**Fig. 3**). The presence of MSFs flanking TF binding sites suggest a regulatory role for these nucleosomes regulating appropriate TF binding. This observation is concordant with similar results from plants and yeast (Zentner and Henikoff 2012, Vera et al., 2014, Pass et al., 2017). These results are consistent with a model in which nucleosomes that provide access to regulatory factor binding sites necessary for a specific genomic response will be more mobile and this will be reflected in greater sensitivity to digestion by MNase.

**Figure 3.**
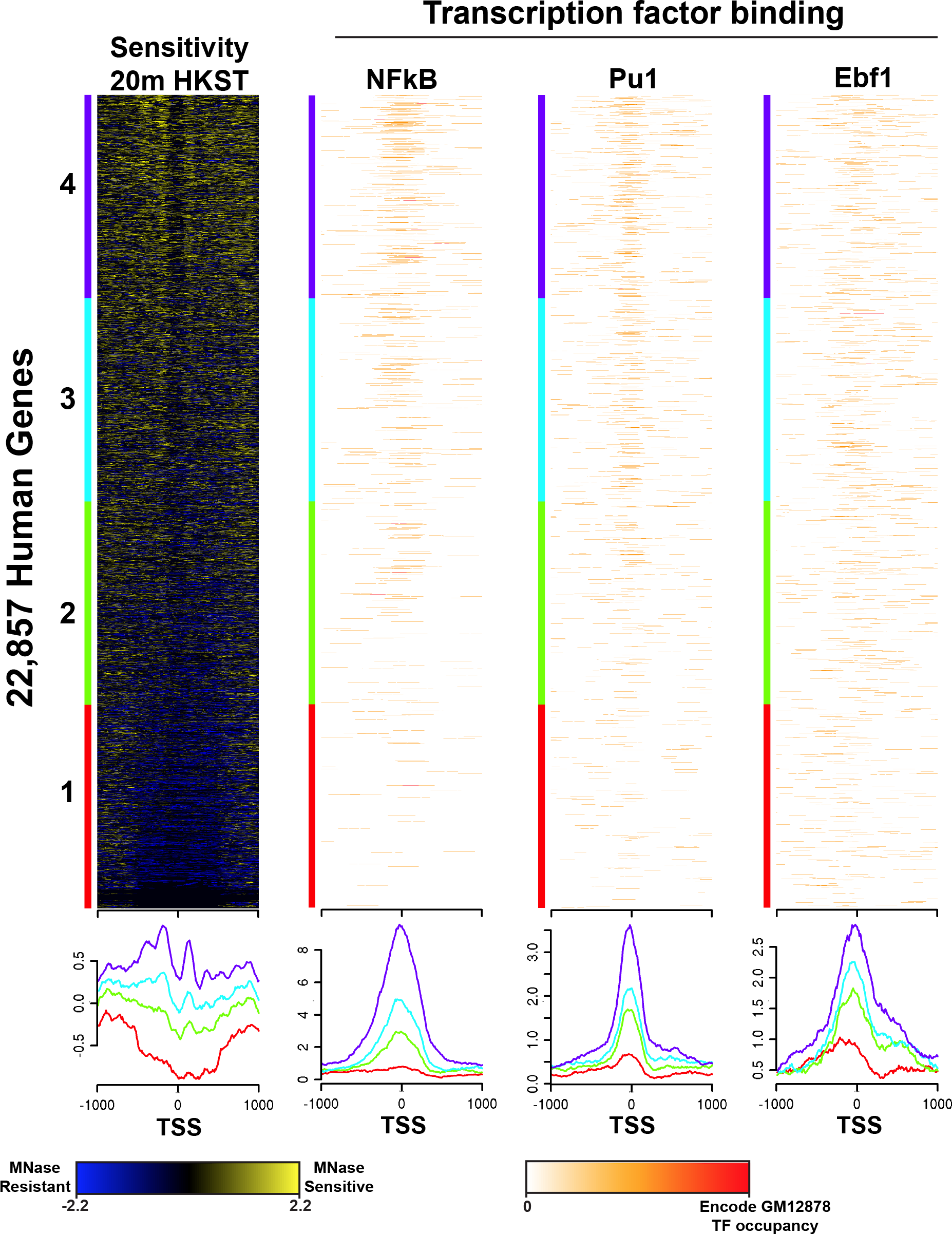
Sensitive nucleosomes flank transcription factor binding sites. MNase-sensitivity of nucleosomes at 20m post-HKST stimulation was sorted into quartiles based on maximum sensitivity followed by ChIPseq peaks of immune transcription factors NFkB, Pu1, Ebf1 in the same sort order.

## Discussion

In this study we measured changes in chromatin structure at promoters in the human lymphoblastoid cell line GM12878 in response to stimulation with HKST. Nucleosomal dynamics occur on a time-scale commensurate with known signaling cascades (Dal Porto et al., 2001, Harwood and Batista, 2008, Arpaia et al., 2011, Browne 2012). Specifically, we observed an increase of a −1 positioned nucleosome at more than 2,000 promoters 20 minutes after stimulation with HKST, and these loci represent 14.7% of the top quartile of expressed genes. This is consistent with important regulatory functions of the −1/+1 nucleosomes reported first in yeast and *Drosophila*, such as structure of a nucleosome-free region at the TSS, interaction with RNA Pol II and transcription factor binding and access to underlying DNA (Svaren and Horz 1995, Jiang and Pugh 2009, Radman-Livaja and Rando 2010, Ballare et al. 2013, Nie et al. 2014, Voong et al. 2016). Additionally, 800 of these genes are involved with regulation of RNA pol II transcription in response to stress and recombination of immunoglobulin genes involved in immune response (**Table S1**). These changes are likely the direct result of HKST immune stimulus producing an inflammatory signaling cascade that induces chromatin regulatory mechanisms at the appropriate promoters. We also observe that the −1 nucleosome occupancy at the 40 and 60 minute post-stimulus timepoints begins to return to the untreated chromatin architecture. These results are consistent with the observation that nucleosome remodeling is a transient event (Sexton et al., 2014, Sexton et al., 2016). These results suggest that nucleosome dynamics may be used as a powerful tool to understand the potential of a cell, beyond the that is garnered from gene expression.

We have also measured promoter nucleosome sensitivity to MNase, showing that sensitive nucleosomes are associated with active transcription. 20 minutes after HKST stimulation, we observe greater −1/+1 nucleosome sensitivity that flanks a larger nucleosome-free region at the TSS. These results are consistent with observations of nucleosome structure at active promoters; and, the additional information given by MNase-sensitivity reflects the regulatory potential of loci that contain sensitive nucleosomes, as reported in yeast, plants, and drosophila (Vera et al. 2014, Mieczkowski et al., 2016, Pass et al., 2017, Brahma and Henikoff 2019). We demonstrate that immune transcription factors NFkB, Pu1, and Ebf1 are associated with positioned sensitive nucleosomes (Garrett-Sinha et al., 1999, Somasundaram et al., 2015, Schwartz et al., 2016, Willis et al. 2017, Zhang et al., 2017). Time course studies have shown early and late gene expression changes during B-cell activation, commensurate with the chromatin structure changes found at promoters following HKST treatment (Fowler et al., 2015, Hawkins et al. 2015). These changes represent a new biochemical potential for cells, and studies of the misregulation of the IKK/NFkB pathway in lymphomas and leukemias show that signal transduction occurs as quickly as 15 minutes with TNFalpha stimulation (Staudt 2010, Hsieh and Van Etten 2014). *In vitro* studies have shown that TF binding to nucleosomal DNA requires nucleosome eviction or repositioning, however pioneer TFs can bind to DNA within the bounds of the nucleosome (Zaret and Carroll 2011, Iwafuchi-Doi and Zaret 2014, Zhu et al. 2018). NFkB has been studied in depth regarding its ability to bindchromatin and induce immune gene transcription, and it appears to play a diverse role in promoter binding and activation of genes in hetero- and euchromatic regions of the genome (Lone et al. 2013, Bhatt and Ghosh 2014, Cieślik and Bekiranov 2015). The association of these immune TFs with increased sensitive nucleosomes at the 20 minute time point suggests that changes in sensitivity create a new chromatin landscape at appropriate promoters, potentiating regulatory factor binding.

A genomic response occurs through multiple regulatory layers, including signaling cascades, regulatory factors, and the resulting regulation of chromatin structure and gene expression. Our study adds to a comprehensive understanding of the induction of immune signaling pathways. Our results add important and complementary information to existing data showing the importance of the chromatin landscape at promoters including nucleosome position, localization of regulatory factors and histone post-translational modifications. The data we show here expounds upon the interplay between chromatin structure to function, which is consonant with models suggested by preeminent immunologists (Natoli 2010, Cuartero et al., 2018, Zhang and Cao 2019). In aggregate, our results suggest that chromatin structure plays a functional role in the dynamic immune response and that nucleosome sensitivity indicates the regulatory potential of specific loci that are poised for the appropriate genomic response.

## Supplementary Information

**Figure S1.**
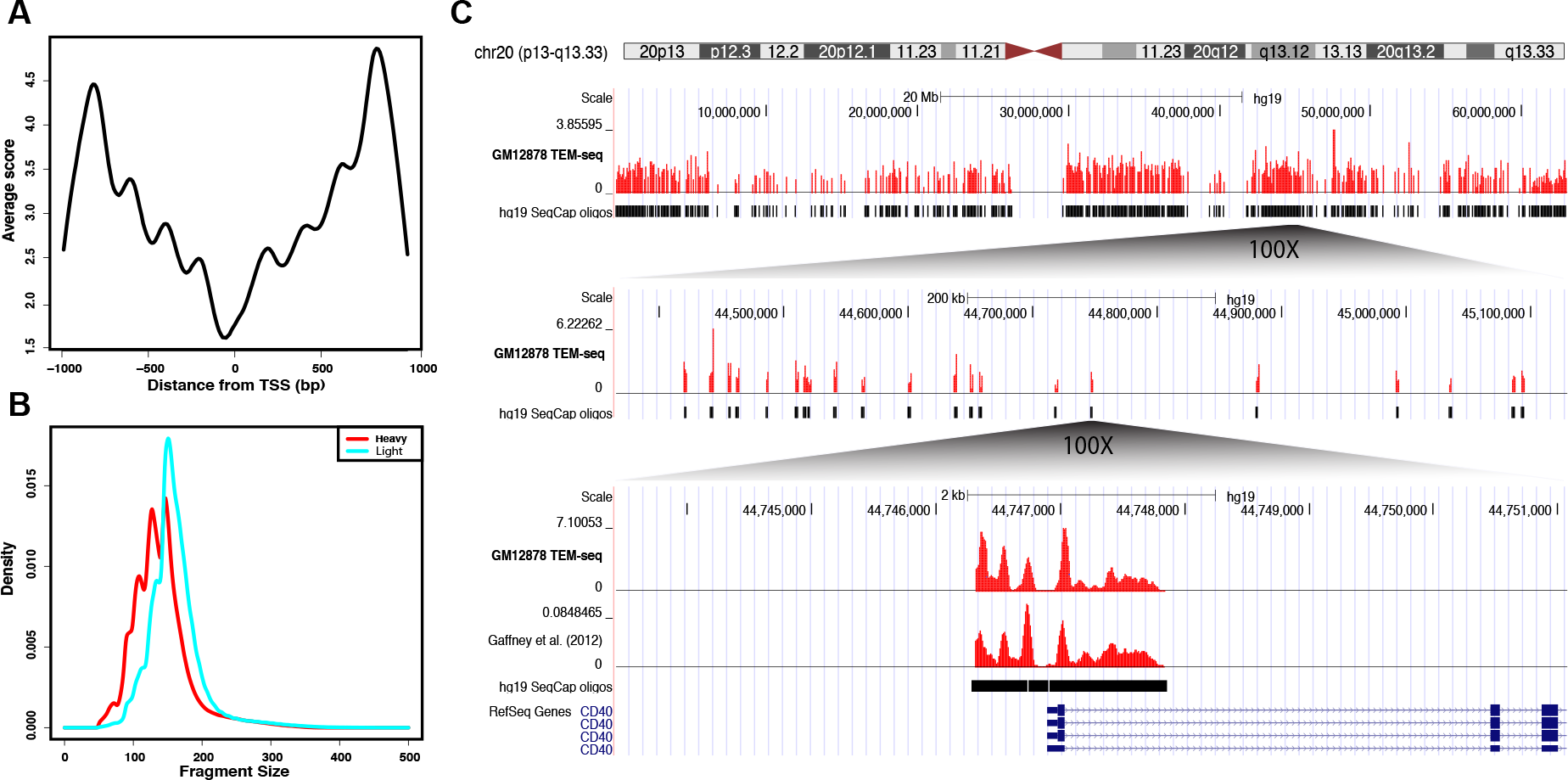
Specific enrichment of MNase-seq fragments at promoters using mTSS-Seq, related to all figures. (A) Average plot of sequence-captured GM12878 MNase-seq data across the TSS +/− 1000bp, all fragments. (B) Histogram of fragment sizes for GM12878 heavy and light digests (C) Schematic demonstrating targeted enrichment with hg19 SeqCap oligos (mTSS-seq). GM12878 Untreated data for chromosome 20 shown in the UCSC genome browser (genome.ucsc.edu), with multiple magnifications down to a single TSS, CD40, compared to a previously published nucleosome mapping data set from Gaffney et al., 2012.

**Figure S2.**
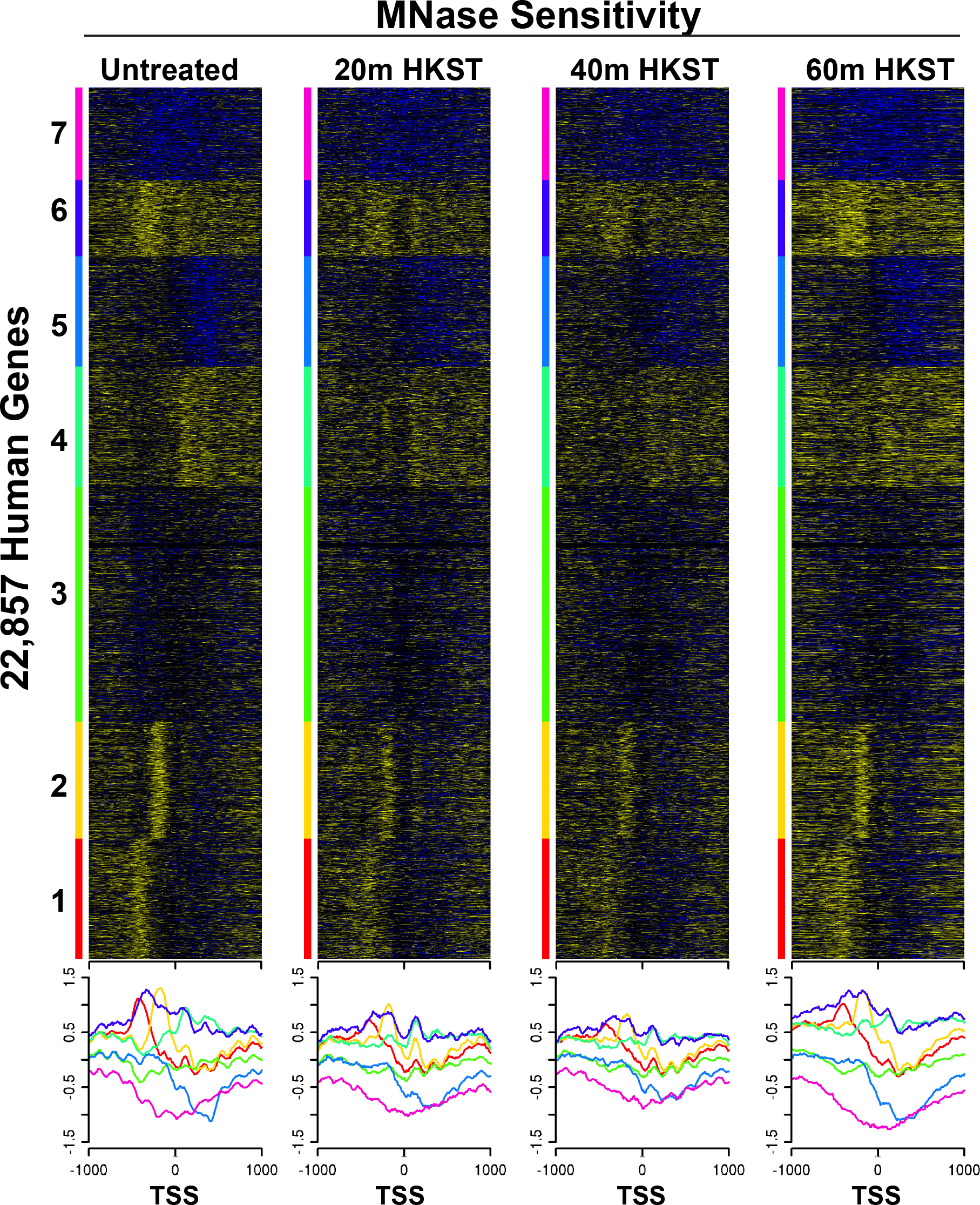
MNase-sensitive of all promoters based on untreated GM12878 cells, related to Figure 2. The log_2_ratio of light/heavy MNase digest (MNase sensitivity) was determined for all human promoters centered on the TSS (+/−1kb). The MNase sensitivity for the untreated control cells was sorted with k-means clustering (k=7) followed by the MNase sensitivity for the 20, 40, and 60 minute post-HKST stimulation data sets on the same sort order. Average profiles per cluster are shown below the heatmaps for each time point. The left most heatmap dictates the sort order for all other maps in the panel.

**Table S1. Results of the gene ontology analysis and gene lists for cluster 5, related to Figure 1**.

**Table S2. Results of the gene ontology analysis and gene list for cluster 1, related to Figure 1**.

